# Multiregional blood-brain barrier phenotyping identifies the prefrontal cortex as the most vulnerable region to ageing in mice

**DOI:** 10.1101/2025.03.09.642233

**Authors:** Isabel Bravo-Ferrer, Katrine Gaasdal-Bech, Chiara Colvin, Hollie J. Vaughan, Jonathan Moss, Anna Williams, Blanca Díaz Castro

**Author notes:** Isabel Bravo-Ferrer^£^ - Department of Cell and Developmental Biology, Division of Biosciences, University College London, London, United Kingdom. Hollie Vaughan -Centre for Inflammation Research, Institute for Regeneration and Repair, University of Edinburgh, Edinburgh, United Kingdom. Jonathan Moss - Roslin Institute 3D Volume Electron Microscopy Capability, Proteomics and Metabolomics Facility, The Roslin Institute, Royal (Dick) School of Veterinary Studies, College of Medicine and Veterinary Medicine, University of Edinburgh, Scotland, UK. Equally contributing authors (Isabel Bravo-Ferrer, Katrine Gaasdal-Bech).

## Abstract

Age-associated vascular alterations make the brain more vulnerable to neuropathologies. Research in humans and rodents have demonstrated structural, molecular, and functional alterations of the aged brain vasculature that suggest blood-brain barrier (BBB) dysfunction. However, these studies focused on particular features of the BBB and specific brain regions. Thus, it remains unclear if and which BBB age-associated phenotypes are conserved across brain areas. Moreover, there is very limited information about how BBB dysfunction and cell-specific phenotypes relate to each other. In this manuscript, we use immunofluorescence, transmission electron microscopy (TEM), and permeability assays to assess how age-associated BBB molecular, structural, and functional phenotypes correlate between the BBB cell types at three brain regions (prefrontal cortex, hippocampus, and corpus callosum) during mouse early ageing. We discovered that at 18-20 months of age, the mouse prefrontal cortex BBB is the most affected region, with alterations in brain endothelial cell protein expression, BBB permeability, basement membrane thickness, and astrocyte endfoot size when compared to young mice. Here, we deliver a detailed multicellular characterisation of region-dependent BBB changes at early stages of ageing. Our data paves the way for future studies to investigate how region-specific BBB dysfunction may contribute to disease-associated regional vulnerability.

## Introduction

Age-associated alterations make the brain more vulnerable to neuropathologies ^1^. In the brain, vascular dysfunction appears as a prominent feature of age that often precedes cognitive decline ^2–4^. Research in humans and rodents have demonstrated structural ^5–7^, molecular ^8–10^, and functional ^11–13^ alterations of the brain vasculature with age that suggest blood-brain barrier (BBB) dysfunction.

The BBB is essential to maintain a healthy brain microenvironment as it strictly regulates the exchange of molecules between the blood and the brain. In the capillaries, constituting 85% of the brain vasculature, the BBB is formed by the brain endothelial cells (BECs), pericytes, basement membrane, and astrocyte endfeet, which are specialised perivascular subcellular compartments. The BECs form the vessel wall and are the first barrier of the BBB. They have a luminal side facing the blood and an abluminal side that is surrounded by the basement membrane and faces pericytes and astrocyte endfeet ^14^. To form an effective BBB, BECs restrict entry by expressing robust molecular complexes called tight junctions, which firmly bind BECs, and by maintaining a low transcytosis rate, limiting the movement of non-regulated cargo ^15^. The import of necessary substances to the brain side is ensured by molecule-specific transmembrane transporters ^16^. Embracing the BECs, pericytes enhance their BBB properties and regulate blood flow ^17^. Lastly, astrocytes uniquely enwrap the brain vasculature while simultaneously contact brain cells to coordinate vascular and neural functions ^17–20^. For example, astrocytes take up nutrients transported in the blood and metabolise them ^21,22^, modulate local blood flow ^23–26^, and enable the clearance of brain by-products and toxins ^27,28^.

With age, the BBB deteriorates in both mouse and humans. In BECs, the expression of tight junctions has been shown to be reduced ^5,7,8^, transcytosis increased ^12^, and vesicular and molecule-specific transporters altered ^5,10^. In addition, a reduction of pericyte vessel coverage has been reported in aged brains ^12,29,30^, while the basement membrane, the vascular extracellular matrix, increases in size ^5–7,9^. Astrocyte endfeet have also been reported to be affected by age, with alterations in the cellular polarisation of the astrocyte water channel aquaporin-4 (AQP-4) in mouse and human ^11,31^ and hypertrophic appearance in the brains of aged rats ^5^.

Overall, numerous studies have demonstrated that age affects the BBB function, structure, and molecular profiles in both rodents and humans. However, these studies focused on particular aspects of the BBB and specific brain regions. Thus, important questions emerge. Are BBB age-associated phenotypes conserved across brain regions or are they region-specific? Is there a region-dependent relationship between changes observed in each component of the BBB?

In this manuscript, we use immunofluorescence, transmission electron microscopy (TEM), and cadaverine permeability assays to assess how age-associated BBB molecular, structural, and functional phenotypes correlate between cell types and brain regions during mouse ageing. We pay particular attention to the BECs, which form the main barrier, and the astrocyte endfeet, highly understudied in this context, despite their essential roles as mediators of vascular and neural functions. Moreover, we make side-by-side comparisons of three brain regions of high relevance to neurological conditions that lead to cognitive decline – prefrontal cortex, hippocampus, and corpus callosum, in both male and female mice. Our data paves the way for future studies to investigate how BBB dysfunction may contribute to disease-associated regional vulnerability.

## Results

### Region-specific effects of ageing on BEC features

We assessed BEC phenotype with age, in young (2-4 months) and aged (18-20 months) mice, across the prefrontal cortex, hippocampus, and corpus callosum.

As tight junctions are an essential feature of BECs, we quantified the tight junction protein claudin-5 by immunofluorescence and tight junction tortuosity ^32^ with TEM in young and aged mouse brain tissue. We found no age-associated differences in these measures in any of the three brain regions of study (Fig. 1A-B).

**Figure 1.**
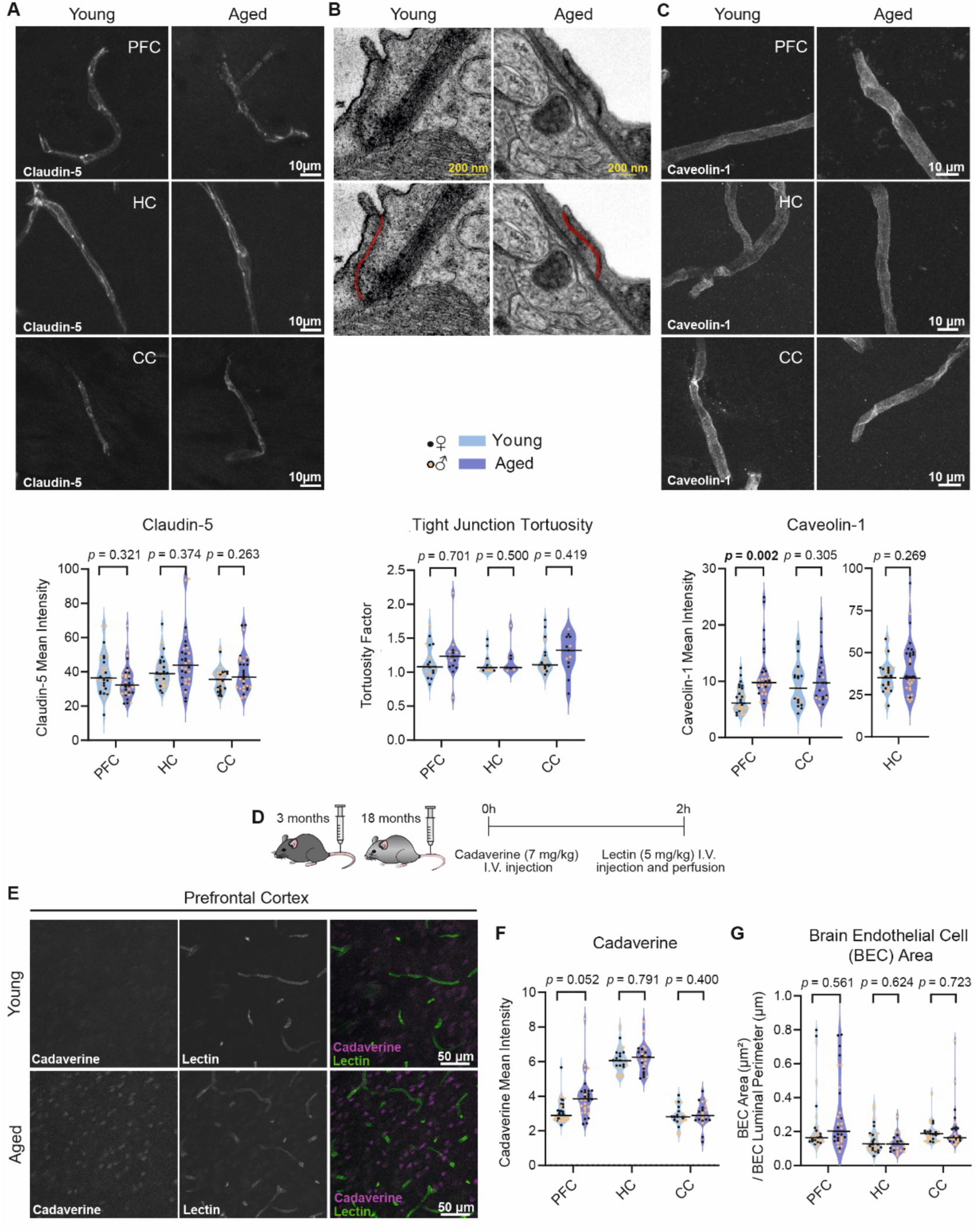
Region-specific effects of ageing on BEC features. A: Representative images of claudin-5 immunostaining in the prefrontal cortex (PFC), hippocampus (HC), and corpus callosum (CC) of young and aged mice with corresponding quantification of staining intensity in individual vessels below. Prefrontal cortex and hippocampus: n=32 vessels from 8 mice (4 females, 4 males). Corpus callosum: n=32 vessels from 8 mice (4 females, 4 males). B: Representative images of prefrontal cortex tight junctions (coloured in red) of young and aged mice with quantification of tight junction tortuosity in individual vessels in the prefrontal cortex (PFC), hippocampus (HC) and corpus callosum (CC) of young and aged mice below. PFC: n=15-19 vessels from 6 mice (3 females, 3 males). HC: 8-13 vessels from 5 mice (young: 3 females, 2 males; aged: 2 females, 3 males). CC: 13-20 vessels from 6 mice (3 females, 3 males). C: Representative images of caveolin-1 immunostaining in the prefrontal cortex (PFC), hippocampus (HC) and corpus callosum (CC) of young and aged mice with corresponding quantification of staining intensity in individual vessels below. PFC and HC: n=32 vessels from 8 mice (4 females, 4 males). CC: n=24 vessels from 8 mice (4 females, 4 males). D: Schematic representing the experimental design to determine the BBB extravasation of cadaverine in young and aged mice. E: Representative images of cadaverine signal in the prefrontal cortex of young and aged mice. F: Quantification of cadaverine mean intensity in the prefrontal cortex (PFC), in the hippocampus (HC), and in the corpus callosum (CC). PFC: n=32 images from 8 mice (4 females, 4 males). HC: n=24 images from 8 mice (4 females, 4 males). CC: n=24 images from 8 mice (4 females, 4 males). G: Quantification of BEC area in individual vessels in the prefrontal cortex (PFC), hippocampus (HC) and corpus callosum (CC) of young and aged mice. PFC and CC: n=30 vessels from 6 mice (3 females, 3 males). HC: 25-30 vessels from 5-6 mice (young: 3 females, 3 males; aged: 2 females, 3 males). See Table 3 for further description on each measurement. Data were analysed using linear mixed-effects modelling (LMM; Intensity ∼ Group + (1 | Animal) + (1 | Sex)) followed by Type 3 ANOVA with Satterthwaite approximation. The horizontal bars on the violin plots represent the median.

BECs are characteristic for their low levels of caveolae-mediated transcytosis ^33^ which can be altered by age ^12^. Thus, we investigated changes in expression of caveolin-1, the principal component of caveolae ^34^, in BECs by immunofluorescence. We observed increased expression of caveolin-1 in the aged prefrontal cortex when compared to their young counterparts, with no differences in the aged hippocampus or corpus callosum (Fig. 1C). Transcellular transport enhancement can lead to BBB permeability increase ^12,35^, so we next evaluated permeability changes associated to caveolin-1 upregulation by quantifying the extravasation of a small dye that does not cross the BBB under healthy conditions (cadaverine-546 of 800 Da) (Fig. 1D). We observed a subtle increase (*p* = 0.052) of cadaverine transport across the BBB in the prefrontal cortex of aged compared to young mice, not apparent in hippocampus or corpus callosum (Fig. 1E-F and Supplementary Fig. 1). There was no age-related BEC size alteration observed with TEM in any of the three brain regions of interest (Fig. 1G and Supplementary Fig. 4-6).

Our data suggests that with age there is increased caveolin-1 mediated transcellular permeability confined to the prefrontal cortex (compared to hippocampus and corpus callosum) whilst other BEC features like tight junctions or size remain spared.

### Basement membrane age-associated alterations

Using TEM, we also assessed basement membrane thickness, which plays a critical role as an interface for cell-cell communication and which has been previously shown to increase with age ^5–7,9^. Unlike previous studies, in our analysis we distinguished the basement membrane segments located between BECs and pericytes from those between BECs and astrocyte endfeet to assess if they were differentially altered. Consistent with earlier findings, we observed an increase in basement membrane thickness with age, but only in the prefrontal cortex and only for the basement membrane between astrocyte endfeet and BECs (Fig. 2A, B, Supplementary Fig. 2A and 4-6). This result, together with the increase in caveolin-1 expression (Fig. 1A), suggested higher BBB susceptibility to early ageing in the prefrontal cortex when compared to hippocampus and corpus callosum, and it prompted us to investigate if these alterations may coexist with changes in astrocyte endfeet.

**Figure 2.**
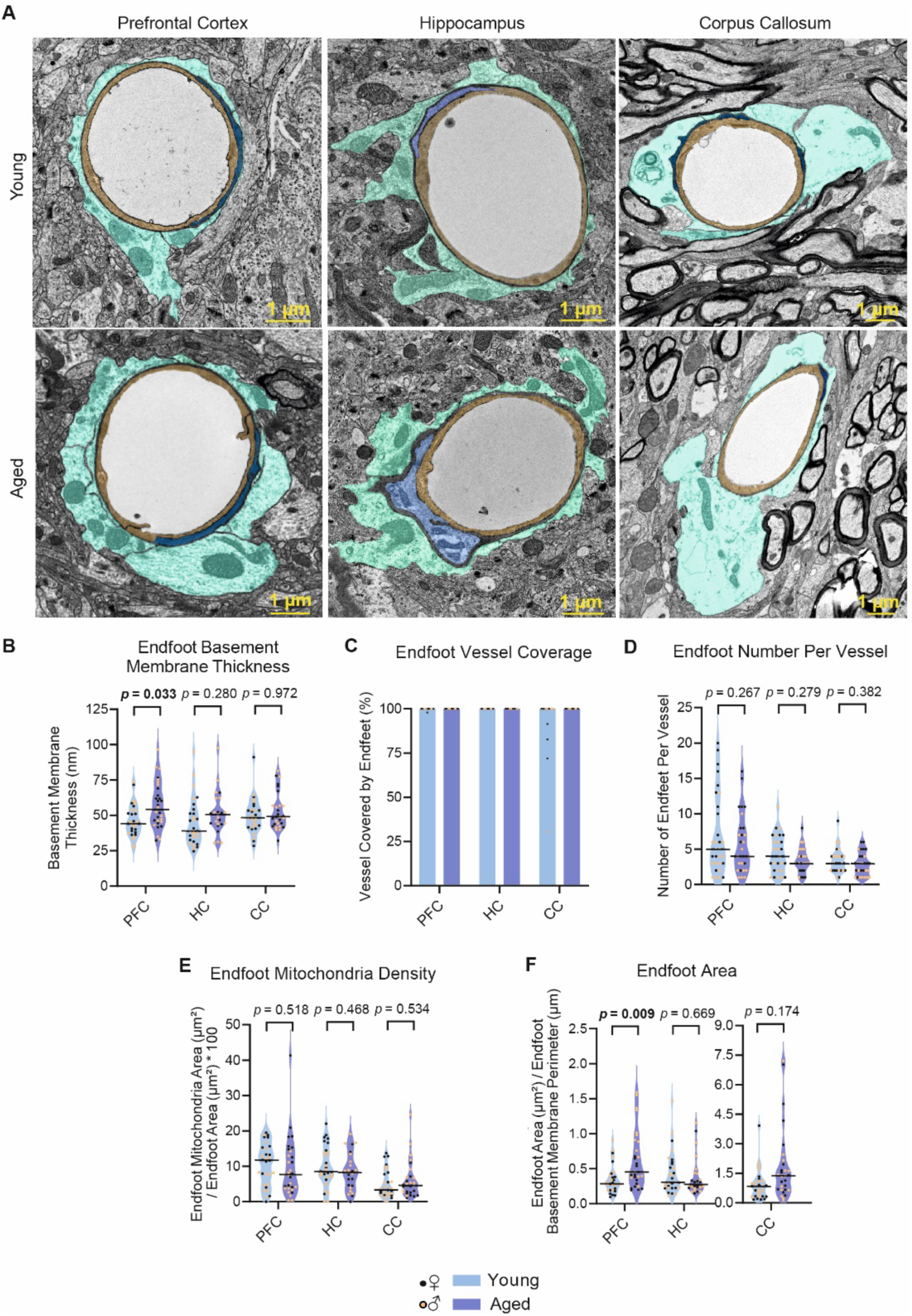
Region-specific effects of ageing in astrocyte endfoot ultrastructural features. A: Representative images of blood vessels in the prefrontal cortex (PFC), hippocampus (HC) and corpus callosum (CC) of young and aged mice. BECs are coloured in yellow, pericytes are coloured in blue and endfeet are coloured in cyan. B-F: Quantification of endfoot basement membrane thickness (B), endfoot vessel coverage (C), endfoot number per vessel (D), endfoot mitochondria density (E), and endfoot area (F) in individual vessels in the PFC, HC and CC of young and aged mice. See Table 3 for further description on each measurement. PFC and CC: n=30 vessels from 6 mice (3 females, 3 males). HC: 25-30 vessels from 5-6 mice (young: 3 females, 3 males; aged: 2 females, 3 males). Data were analysed using LMM (Intensity ∼ Group + (1 | Animal) + (1 | Sex)) followed by Type 3 ANOVA with Satterthwaite approximation. The horizontal bars on the violin plots represent the median. Figure 4G does not display statistical analysis because, for most of the conditions, astrocyte coverage was 100% in all vessels and this led to lack of variance in the dataset which made it unsuitable for statistical analysis. In this figure, some points in young CC have a value <100%; this was due to the presence of ambiguous dark endfoot-like structures that were not included in the counting. Only clear endfeet were considered because we could not determine the cell identity of the dark structures.

### Region-specific effects of ageing in astrocyte endfeet

We began by evaluating molecular changes in astrocyte endfeet by assessing the expression levels of well-known endfoot proteins: aquaporin-4 (AQP-4), β-dystroglycan, and dystrophin by immunofluorescence. AQP-4 is a water-selective channel with proposed roles in brain water homeostasis and glymphatic flow ^36,37^. We calculated the polarisation ratio of vascular versus parenchymal AQP-4 expression in prefrontal cortex, hippocampus, and corpus callosum and identified no age-associated differences (Supplementary Fig. 3A-C, 3H-J, and 3O-Q). We also examined β-dystroglycan and dystrophin, two components of the dystrophin-associated protein complex (DAPC) that anchors AQP-4 to the plasma membrane of the astrocyte endfoot ^18,38^. Although there was no change in these proteins with age in the prefrontal cortex or corpus callosum (Supplementary Fig. 3D-G and 3R-U), hippocampal β-dystroglycan was significantly higher in aged vessels when compared to the young ones (Supplementary Fig. 3 K-N).

We next used TEM to investigate other, structural characteristic features of astrocyte endfeet. Endfoot coverage and the number of endfeet surrounding each vessel, as well as mitochondrial density displayed no differences with age across any of the three brain regions of study (Fig. 2C-E and Supplementary Fig. 4-6). Despite previous reports showing that pericyte coverage is reduced with age using immunohistochemistry ^12,29,30^, we observed no alterations of pericyte coverage or area with age in any of the brain regions analysed using TEM (Supplementary Fig. 2B-C). However, astrocyte endfeet were significantly hypertrophic in the prefrontal cortex of aged mice compared to young ones, with no differences in hippocampus or corpus callosum (Fig 2F and Supplementary Fig. 4-6).

In summary, we observed that at early stages of ageing, in 18-20-month-old mice, changes to the BBB are subtle, with most of the BBB age-dependent changes presenting in the prefrontal cortex (Table 1).

**Table 1:**
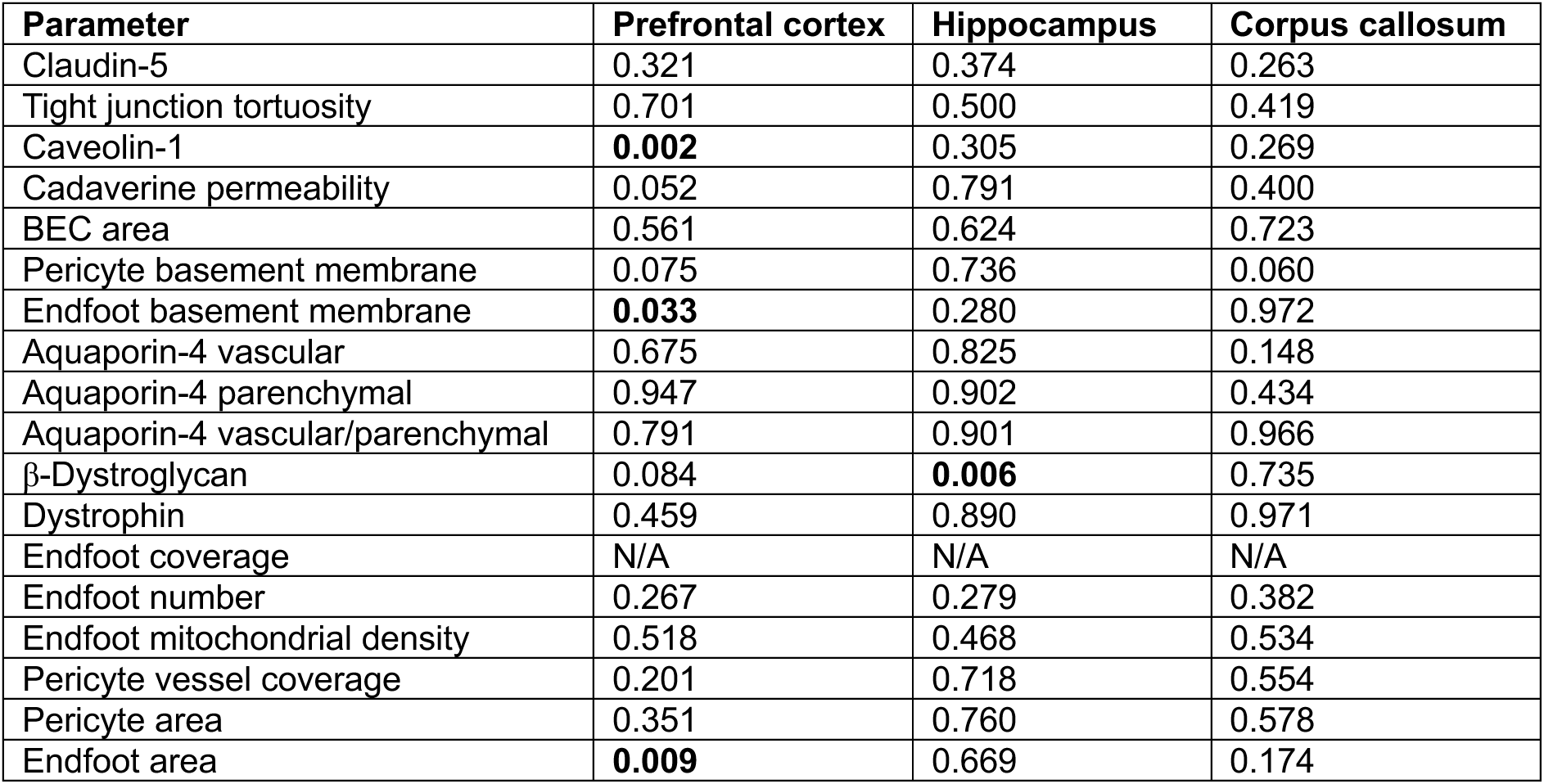
Compilation of *p* values obtained from all the statistical comparisons between aged and young animals in this manuscript.

| Parameter | Prefrontal cortex | Hippocampus | Corpus callosum |
| --- | --- | --- | --- |
| Claudin-5 | 0.321 | 0.374 | 0.263 |
| Tight junction tortuosity | 0.701 | 0.500 | 0.419 |
| Caveolin-1 | **0.002** | 0.305 | 0.269 |
| Cadaverine permeability | 0.052 | 0.791 | 0.400 |
| BEC area | 0.561 | 0.624 | 0.723 |
| Pericyte basement membrane | 0.075 | 0.736 | 0.060 |
| Endfoot basement membrane | **0.033** | 0.280 | 0.972 |
| Aquaporin-4 vascular | 0.675 | 0.825 | 0.148 |
| Aquaporin-4 parenchymal | 0.947 | 0.902 | 0.434 |
| Aquaporin-4 vascular/parenchymal | 0.791 | 0.901 | 0.966 |
| β-Dystroglycan | 0.084 | **0.006** | 0.735 |
| Dystrophin | 0.459 | 0.890 | 0.971 |
| Endfoot coverage | N/A | N/A | N/A |
| Endfoot number | 0.267 | 0.279 | 0.382 |
| Endfoot mitochondrial density | 0.518 | 0.468 | 0.534 |
| Pericyte vessel coverage | 0.201 | 0.718 | 0.554 |
| Pericyte area | 0.351 | 0.760 | 0.578 |
| Endfoot area | **0.009** | 0.669 | 0.174 |

## Discussion

Here we aimed to determine BBB age-related alterations between three brain regions that are highly relevant to cognition: the prefrontal cortex, hippocampus, and the corpus callosum. We discovered that the most vulnerable area to early BBB ageing is the prefrontal cortex with increased caveolae-mediated transcytosis, basement membrane thickness, and endfoot size (Table 1). The hippocampal BBB only presented higher expression of β-dystroglycan in astrocyte endfeet with age (Table 1), and surprisingly, despite the high sensitivity of white matter to age-associate cerebral small vessel disease ^39^, we observed no differences in the corpus callosum BBB (Table 1).

It has been suggested that BEC alterations with age lead to failure in both paracellular ^7,8,40^ and transcellular ^12^ barrier mechanisms, with the observation of increased age-associated BBB permeability in both mouse and human in some studies ^12,13,30,40–42^, but not in others ^8,43^. Whilst altered expression or re-arrangement of tight junction proteins like claudin-5 and ZO-1 have been reported in the brains of aged mice and humans with immunostaining and TEM ^7,8,40^, our study found neither. This discrepancy may be due to differences in the regions of study, ages, and the techniques used. For example, Frias-Anaya et. al. observed that 3D-TEM is needed to be able to detect alterations of the tight junction structure with age in 24-month-old mice ^7^, and only observed alterations with age in the cortex and not the hippocampus. We observed a subtle increase in BBB permeability in the prefrontal cortex (Fig. 1E, 1H) along with increased expression of caveolin-1 with age (Fig. 1A-C), and this was consistent with previously described age-associated increased caveolae-mediated transcytosis ^12^.

Increased basement membrane area in the rodent cortex, hippocampus, thalamus, and striatum has been associated with age in numerous TEM studies ^5–7,9^. In our analysis, we distinguished between the pericyte and endfoot basement membrane and found the endfoot basement membrane was significantly thicker in the 18-20 month aged prefrontal cortex (Fig. 2B) but not in hippocampus or corpus callosum. Why the basement membrane increases in size with age is not known; but previous studies have shown alterations in its protein composition in cortex and hippocampus with age ^9,42,44^.

In line with an increased BBB susceptibility to ageing in the prefrontal cortex, we identified larger astrocyte endfeet in this region in aged mice when compared to the young counterparts, again not observed in hippocampus or corpus callosum endfeet. Hypertrophic endfeet have previously been observed in the striatum of aged rats ^5^. A possibility is that these endfeet are filled with water, although appearance of their cytoplasm was very similar to the one of young endfeet (Supplementary Fig. 4). Whilst, some studies have reported age-associated AQP-4 altered expression in cortical astrocytes ^11,45,46^, we found no AQP-4 expression differences in any region (Supplementary Fig. 3). AQP-4 function can be regulated by its removal from the plasma membrane without the need of relocating within the cell ^47^, a mechanism which would not be detected by our quantification method. Additionally, higher resolution endfoot TEM images could help determine the composition of hypertrophic endfeet, to investigate alternative explanations to an increase in the water content. Interestingly, despite the endfoot change in morphology, neither the number of endfeet nor their vessel coverage was altered. Indeed, virtually complete endfoot coverage of the blood vessels seems to be a very robust feature of the BBB ^14,48^. Mills and colleagues demonstrated that when one endfoot is removed through two-photon ablation, neighbouring astrocytes rapidly extend their processes to replace the ablated structure within minutes even in older mice ^49^.

The limitations of this study include that while we see region-dependent multicellular alterations with age, we do not know if there is a cause-effect among them or if they just represent independent coexisting events. Similarly, we are unclear whether age-driven BBB differences between brain regions relate to parenchymal alterations, or are intrinsic to the vasculature. More comprehensive parallel proteomics and functional studies will give us a more precise picture of the altered pathways that may lead to BBB dysfunction with age, and it would be interesting to include subcortical brain regions that are particularly susceptible to age-related human vascular disease ^39^.

In conclusion, previous research has shown diverse BBB dysfunction phenotypes associated to age in mouse and human. However, the confined focus of each study on either specific brain regions, stages of ageing, or phenotypes has made it difficult to determine if BBB susceptibility in ageing is region-dependent and whether BBB cell-specific alterations correlate with each other. Here, by systematically comparing the phenotypes of the different capillary-BBB components across three brain regions, we demonstrate region-specific changes of the BBB with age. In our study, the hippocampus and corpus callosum appeared resilient to early ageing, while the prefrontal cortex was the most affected region with increased caveolae-mediated transcytosis, basement membrane thickness, and astrocyte endfoot size. New studies will be needed to better understand the mechanisms that lead to region-specific early events of BBB dysfunction, how they increase susceptibility to disease, and if their prevention could delay pathology.

## Materials and methods

### Mice

Animal experiments were conducted in accordance with the national and institutional guidelines ([Scientific Procedures Act] 1986 (UK), and the Council Directive 2010/63EU of the European Parliament and the Council of 22 September 2010 on the protection of animals used for scientific purposes) and had a full Home Office ethical approval. Mice were housed with no restrictions to food and water in a 12-hour light/dark cycle.

All experiments were performed using C57Bl/6J mice (Charles River Laboratories, UK) of 2-3 months or 18-20 months old mice of both sexes.

### BBB permeability assay, vasculature tracing, and drug administration

Alexa 555 conjugated cadaverine (Invitrogen Cat#A30677) was dissolved in sterile PBS at 1mg/ml. Mice received an intravenous (i.v.) injection of 7mg/kg 2 hrs before perfusion. To trace the vasculature, mice received an i.v. injection 5 min before perfusion of fluorescein conjugated lectin from Lycopersicon esculentum (Vector Cat#FL-1171) at a concentration of 5 mg/kg.

### Immunofluorescence

For immunofluorescence studies, mice were subjected to terminal anaesthesia and transcardial perfusion with PBS containing 10 units/mL of heparin followed by 10% of formalin. 40 μm coronal sections were prepared using a crysotat (Leica) and kept in 0.05 M PBS, 250 mM sucrose, 7 mM MgCl2 and 50% glycerol at -20°C. To avoid lipofuscin autofluorescence sections were pre-treated with TrueBlack (Biotium, Cat# 23007) 1X in 70% ethanol for 1 minute before staining. Sections were then washed three times in PBS for 10 min and incubated at room temperature for 2 hr in blocking solution containing 10% of normal goat or donkey serum in PBS with 0.2 Triton X-100 with agitation. Sections were subsequently incubated in primary antibodies diluted in the blocking solution. The following primary antibodies were used: rabbit anti-NeuN (1:2000, Cell Signalling Cat# 12943S), mouse anti-claudin-5 (1:500; Invitrogen 35-2500 and Invitrogen 352588), rabbit anti-caveolin-1 (1:1000, Cell Signalling Cat# 3267), rabbit anti-Pecam-1 (1:1000, BD Bioscience Cat# 550274), rabbit anti-aquaporin4 (1:1000, Millipore Cat# ab3594), mouse anti-dystrophin (1:500, DSHB, RRID: AB_2618143) and mouse anti-dystroglycan (1:1000, DSHB, RRID: AB_2618140). The next day the sections were washed three times in PBS for 10 min each and then incubated with the secondary antibody diluted in blocking solution. The following secondary antibodies were used: goat anti-mouse IgG Alexa 647 (1:1000, Invitrogen Cat# A21235), goat anti-rabbit IgG Alexa 647 (1:1000, Invitrogen Cat# A21244), goat anti-rat IgG Alexa 647 (1:1000, Invitrogen Cat# A21247), goat anti-mouse IgG Alexa 488 (1:1000, Invitrogen Cat# A11001), goat anti-rabbit IgG Alexa 488 (1:1000, Invitrogen Cat# A11008), donkey anti-mouse IgG Alexa 488 (1:1000, Invitrogen Cat# A21202) and donkey anti-goat IgG (1:1000, Invitrogen Cat# A21447). The sections were mounted on microscope slides in ProLong Gold antifade reagent.

### Confocal microscopy and image analysis

Fluorescent images were taken on a Zeiss LSM900 confocal laser-scanning microscope. For BBB permeability assessment, 3-5 images were taken in each region using a 25X 0.8 NA (Plan-Apochromat, Zeiss Cat# 420852-9871-000). For BEC and endfoot markers assessment, 3 (CC) or 4 (PFC, HC) capillaries (diameter < 10 μm) were randomly selected in each region. To do this, a rectangle area within each brain region was selected and points were randomly generated. The closest capillary to each point was then imaged using a 63X 1.4 NA oil immersion objective (Plan-Apochromat, Zeiss Cat# 420782-9900-799). For claudin-5, caveolin-1, pecam-1, dystrophin, and β-dystroglycan, images containing the whole capillary in the z-plane were taken. For aqp4 images, stacks of 6 μm were taken in vessels where the lumen was clearly visible. Imageing settings remained unchanged within groups of the same experiment.

Image analysis was conducted using FIJI (ImageJ). For BBB permeability assessment, the cadaverine mean intensity in each image was measured. For the assessment of claudin-5, caveolin-1, pecam-1, dystrophin, and β-dystroglycan expression, the maximum intensity of z-stack projections from the images were created and channels separated. Using the lectin channel, the vessel area was delimited creating an area of interest (ROI) that was used to measure the mean intensity within the capillary area for each marker. For aqp4 expression assessment, a linear ROI of 5 μm was created perpendicular to the vessel containing both vascular and parenchymal aqp4 signal. The intensity profile of the line was then plotted, and the area under the curve for both vascular and parenchymal was calculated.

### Transmission electron microscopy (TEM)

Mice were first perfused with PBS, followed by a fixative solution composed of 4% paraformaldehyde (w/v) and 2% glutaraldehyde (v/v) in 0.1 M phosphate buffer. Brains were then extracted, immersed in the same fixative for 24 hours at 4°C, and sliced in ice-cold PBS into 50 μm sections using a vibratome.

The tissue sections were subsequently treated with 1% osmium tetroxide in 0.1 M phosphate buffer for 30 minutes, followed by a dehydration process through an ascending ethanol series (50%, 70%, 95%, and 100%) and acetone. Once dehydrated, the sections were immersed into resin and left to set overnight at room temperature. Afterwards, the sections were mounted onto microscope slides, covered with coverslips, and cured at 65°C over three days.

ROIs including the corpus callosum, prefrontal cortex, and hippocampus were dissected from the slides, mounted onto plastic blocks, and further sectioned to ultrathin 60 nm sections for electron microscopy. These sections were then contrasted with lead citrate and uranyl acetate and imaged using a JOEL TEM-1400 Plus electron microscope at the Edinburgh Discovery Research Platform for Hidden Cell Biology. Imageing magnifications were chosen to include each vessel along with the adjacent perivascular area for analysis. All vessels analysed were capillaries (diameter < 10 μm).

### TEM analyses

The number of vessels analysed per region are listed in Table 2. Tight junction tortuosity was calculated as the ratio of the tight junction length to the diagonal of the rectangle containing the entire junction ^50^. Normalized brain endothelial cell area was calculated as the brain endothelial cell area divided by its luminal perimeter, while normalised pericyte area was obtained by dividing the pericyte area by the abluminal perimeter of the brain endothelial cell. Similarly, normalised endfoot area was measured as the ratio of the endfoot area to the perimeter of its basement membrane. Endfoot and pericyte coverage was quantified as the length of the endfoot or pericyte basement membrane normalised to the brain endothelial cell abluminal perimeter and expressed as a percentage. Finally, endfoot mitochondria area was analysed measuring the total area occupied by mitochondria within the endfoot and endfoot mitochondria density was expressed as a fraction of the total endfoot area (Table 3).

**Table 2:**
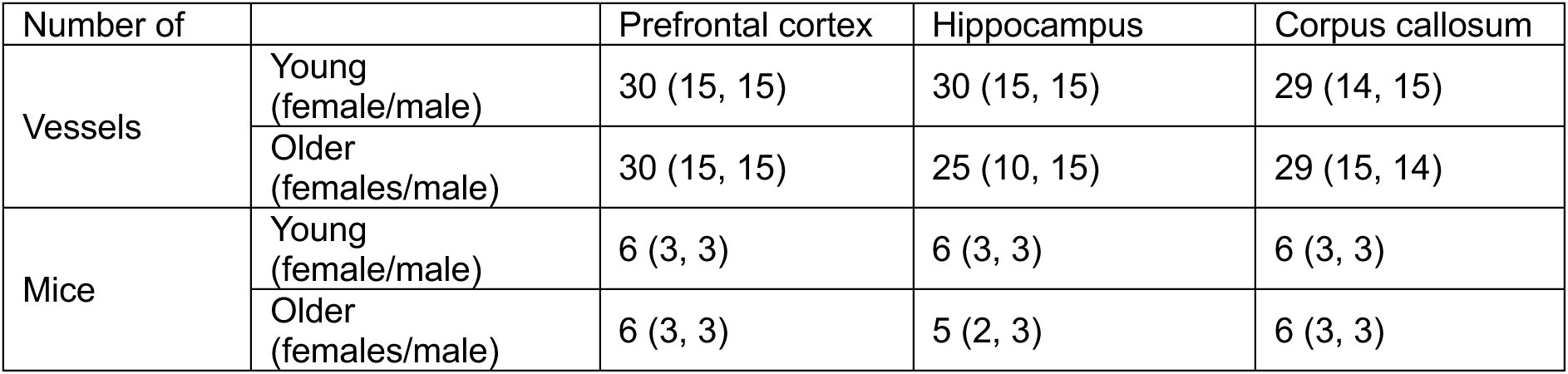
Condition distribution of the images used for the TEM study.

**Table 3:** Summary of measurements taken for the analysis of TEM images.

| Parameter | Measurement |
| --- | --- |
| Tight junction tortuosity | Length of the tight junction / diagonal of the rectangle containing the complete tight junction<sup>51</sup> |
| Normalised brain endothelial cell area | Brain endothelial cell area / brain endothelial cell luminal perimeter |
| Pericyte basement membrane thickness | Pericyte basement membrane area / Pericyte basement membrane length |
| Endfoot basement membrane thickness | Endfoot basement membrane area / Endfoot basement membrane length |
| Endfoot mitochondria density | Endfoot mitochondria area / endfoot area |
| Pericyte coverage | Length of the pericyte basement membrane * 100 / brain endothelial cell abluminal perimeter |
| Endfoot coverage | $\text{Length of the endfoot basement membrane} * 100 / \text{brain endothelial cell abluminal perimeter}$ |
| Normalised pericyte area | $\text{Pericyte area} / \text{brain endothelial cell abluminal perimeter}$ |
| Normalised endfoot area | $\text{Endfoot area} / \text{endfoot basement membrane perimeter}$ |

### Statistics

All analyses were performed blinded to the conditions of study. Data were analysed using GraphPad Prism or R. All comparisons contained more than one datapoint per animal. Thus, to be able to consider multiple measures per animal to take within animal variability into consideration without introducing pseudo-replication, we used linear mixed-effect modelling (LMM) for our analyses (https://github.com/Spires-Jones-Lab/Linear-mixed-models-R). LMM compared between conditions i.e. aged vs young as a fixed effect and included sex and animal as random effects. After assembling an initial LMM, the normality of the residuals was assessed using Shapiro-Wilk test. To meet model assumptions, data with non-normal residuals were transformed using Tukey Ladder of Powers. To identify significance main effects, Type 3 ANOVA with Satterthwaite approximation was run on the LMM. Graph represent all datapoints measured with their median highlighted. Statistical significance level was considered at *p* < 0.05.

## Acknowledgments

This work was supported by UK Dementia Research Institute (UKDRI-4007 and UKDRI-4206), through UK DRI Ltd, principally funded by the Medical Research Council and Alzheimer’s Research UK (ARUK-NC2020-SCO) funds to BDC. KG-B is a student on the Translational Neuroscience PhD Programme and is funded by Wellcome Trust (218493/Z/19/Z). JM and AW were supported by UK Dementia Research Institute (UKDRI–4205). We thank Stephen Mitchell for training and technical expertise with EM at the Discovery Research Platform for Hidden Cell Biology, Valentina Escott-Price for support with statistical assessment and Tara Spires-Jones for sharing the code we used to perform linear mixed-effect model analyses (https://github.com/Spires-Jones-Lab/Linear-mixed-models-R).

## Contributions

IB-F performed most of the experiments and most of the data analyses. KG-B helped with data analysis, made the figures, and performed statistical analyses. CC performed some experiments and analyses. HV performed some analyses. JM and AW advised and supported with electron microscopy expertise that was key to the success of this project. BDC conceived the study, directed the study, and coordinated the collaborative work. BDC and IB-F wrote the paper. All the authors commented on the manuscript.

## Additional information

Supplementary Data (6 figures)

Correspondence to BDC

---

**Supplementary Figure 1.**
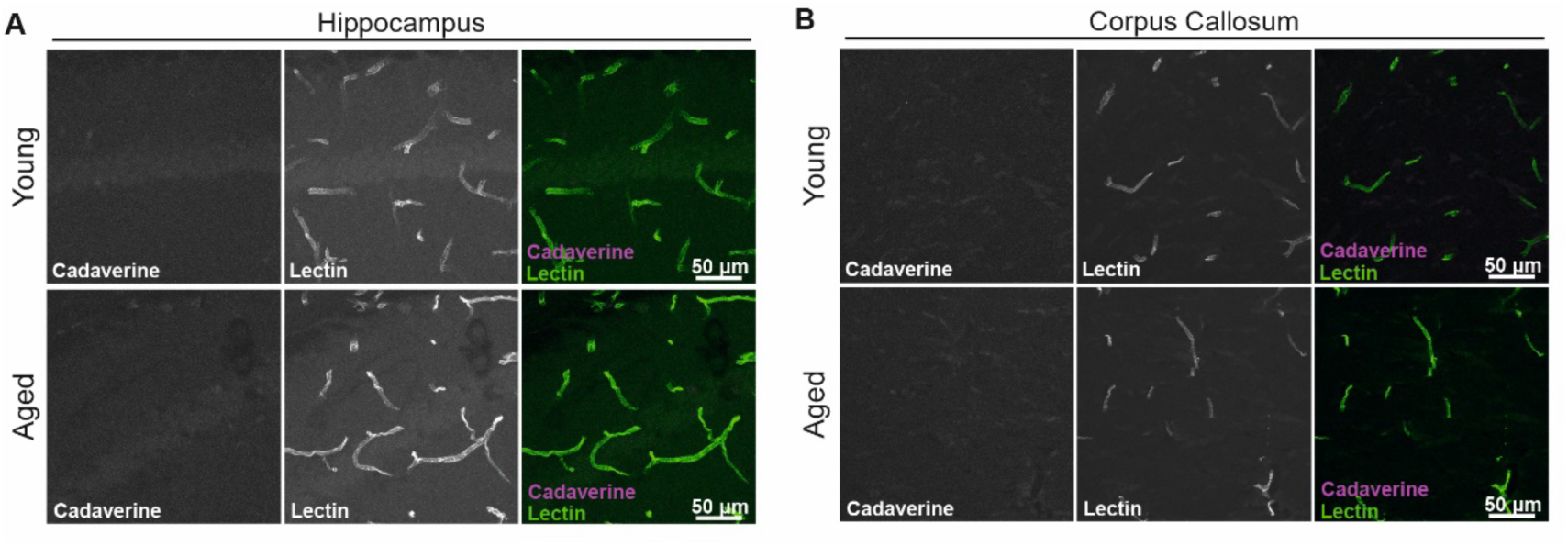
Representative images of cadaverine signal in hippocampus (A) and corpus callosum (B).

**Supplementary Figure 2.**
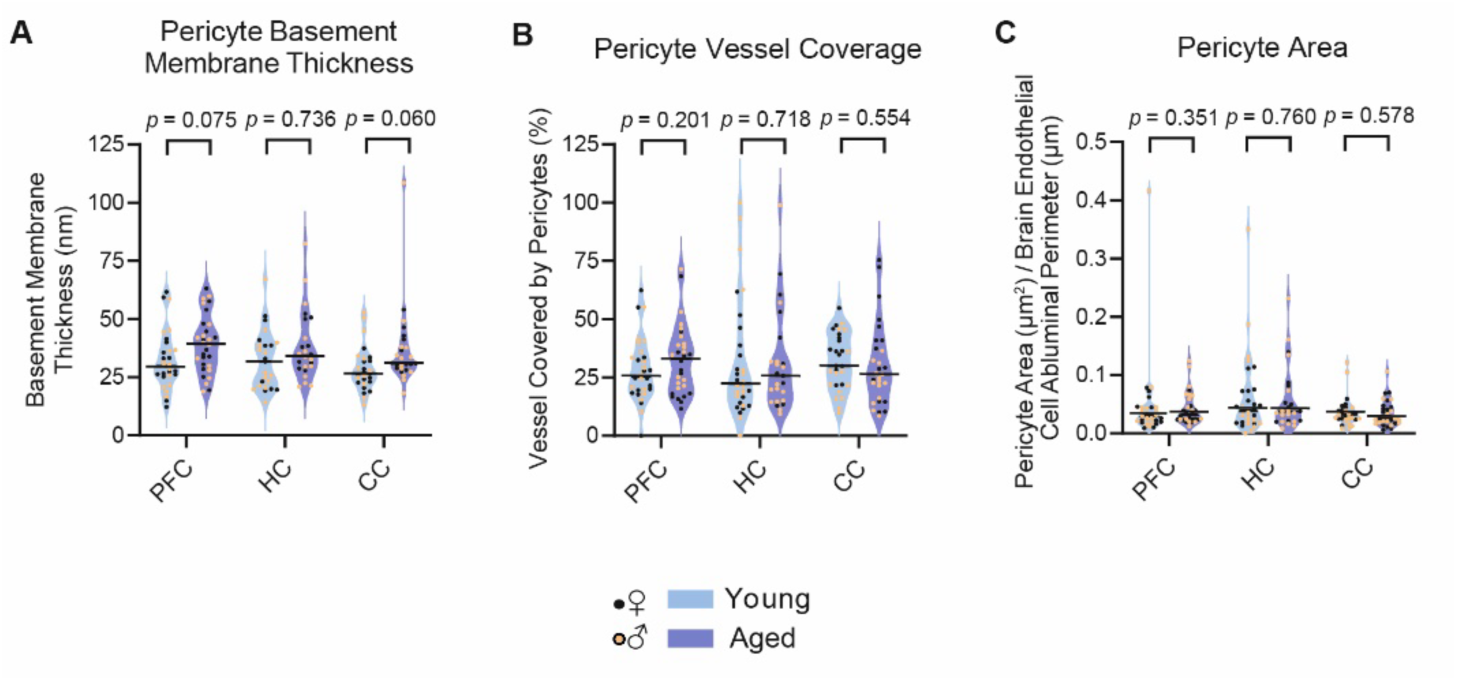
A-C: Quantification of pericyte basement membrane thickness (A), pericyte vessel coverage (B), and pericyte area (C) in individual vessels in the PFC, HC and CC of young and aged mice. See Table 3 for further description on each measurement. PFC and CC: n=30 vessels from 6 mice (3 females, 3 males). HC: 25-30 vessels from 5-6 mice (young: 3 females, 3 males; aged: 2 females, 3 males). Data were analysed using LMM (Intensity ∼ Group + (1 | Animal) + (1 | Sex)) followed by Type 3 ANOVA with Satterthwaite approximation. The horizontal bars on the violin plots represent the median.

**Supplementary Figure 3.**
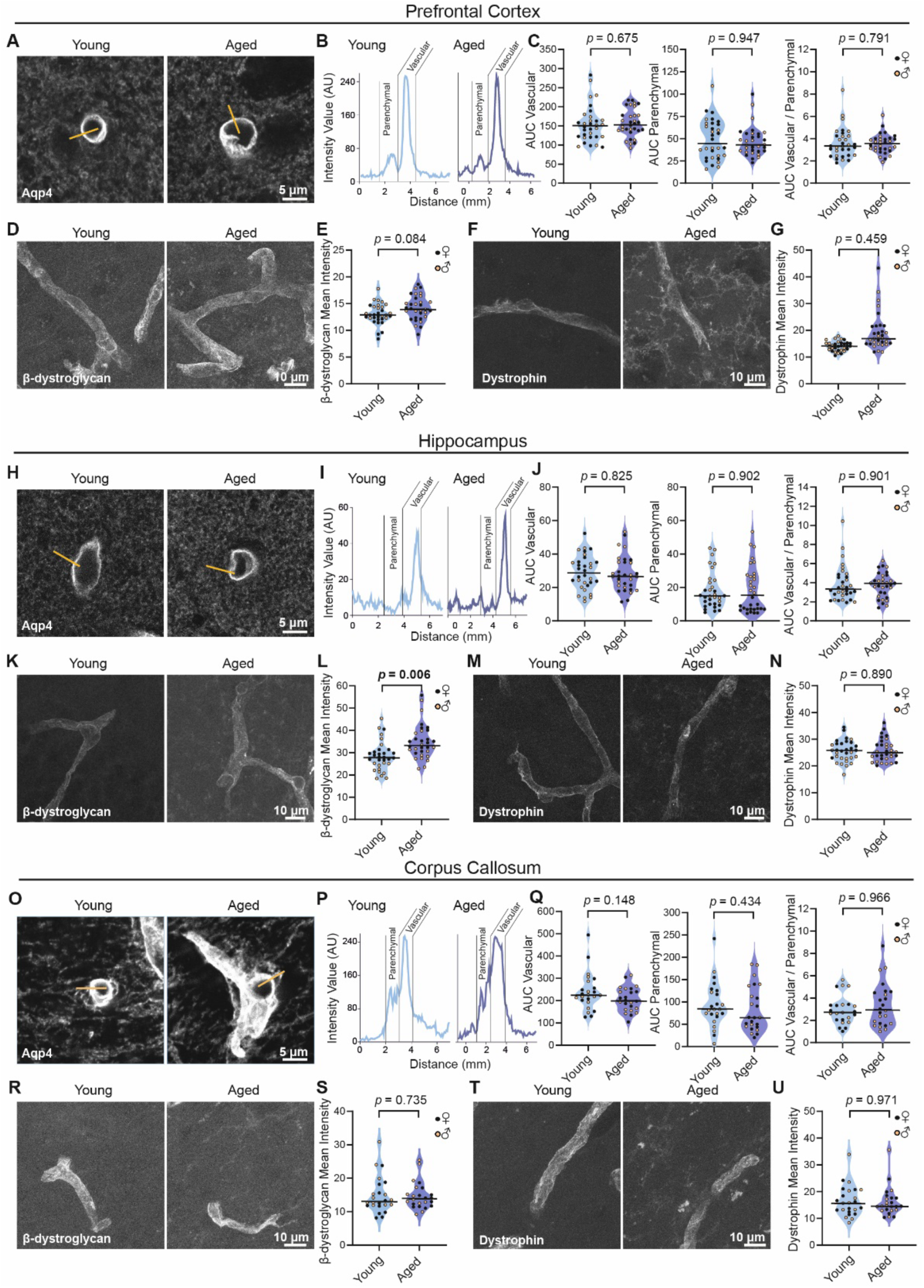
Region-specific effects of ageing in astrocyte endfoot protein expression. A-B: Representative images of AQP-4 immunostaining in the prefrontal cortex of young and aged mice. For the analysis, a yellow line was drawn transversally to individual vessels and the intensity profile along the line was plotted (B), delineating AQP-4 signal across vascular and parenchymal regions. C: Quantification of the area under the curve (AUC) in the vascular (left) and parenchymal (middle) regions and the vascular-to-parenchymal AUC ratio (right) in individual vessels of the prefrontal cortex. D: Representative images of ß-dystroglycan immunostaining in the prefrontal cortex of young and aged mice. E: Quantification of ß-dystroglycan staining intensity in individual vessels of the prefrontal cortex. F: Representative images of dystrophin immunostaining in the prefrontal cortex of young and aged mice. G: Quantification of dystrophin staining intensity in individual vessels of the prefrontal cortex. H: Representative images of AQP-4 immunostaining in the hippocampus of young and aged mice. I: Intensity profiles corresponding to examples shown in H. J: Quantification of the area under the curve (AUC) in the vascular (left) and parenchymal (middle) regions and the vascular-to-parenchymal AUC ratio (right) in individual vessels of the hippocampus. K: Representative images of ß-dystroglycan immunostaining in the hippocampus of young and aged mice. L: Quantification of ß-dystroglycan staining intensity in individual vessels of the hippocampus. M: Representative images of dystrophin immunostaining in the hippocampus of young and aged mice. N: Quantification of dystrophin staining intensity in individual vessels of the hippocampus. O: Representative images of AQP-4 immunostaining in the corpus callosum of young and aged mice. P: Intensity profiles corresponding to examples shown in O. Q: Quantification of the area under the curve (AUC) in the vascular (left) and parenchymal (middle) regions and the vascular-to-parenchymal AUC ratio (right) in individual vessels of the corpus callosum. R: Representative images of ß-dystroglycan immunostaining in the corpus callosum of young and aged mice. S: Quantification of ß-dystroglycan staining intensity in individual vessels of the corpus callosum. T: Representative images of dystrophin immunostaining in the corpus callosum of young and aged mice. U: Quantification of dystrophin staining intensity in individual vessels of the corpus callosum. Prefrontal cortex and hippocampus: n=32 vessels from 8 mice (4 females, 4 males). Corpus callosum: n=24 vessels from 8 mice (4 females, 4 males). Data were analysed using LMM (Intensity ∼ Group + (1 | Animal) + (1 | Sex)) followed by Type 3 ANOVA with Satterthwaite approximation. The horizontal bars on the violin plots represent the median.

**Supplementary Figure 4.**
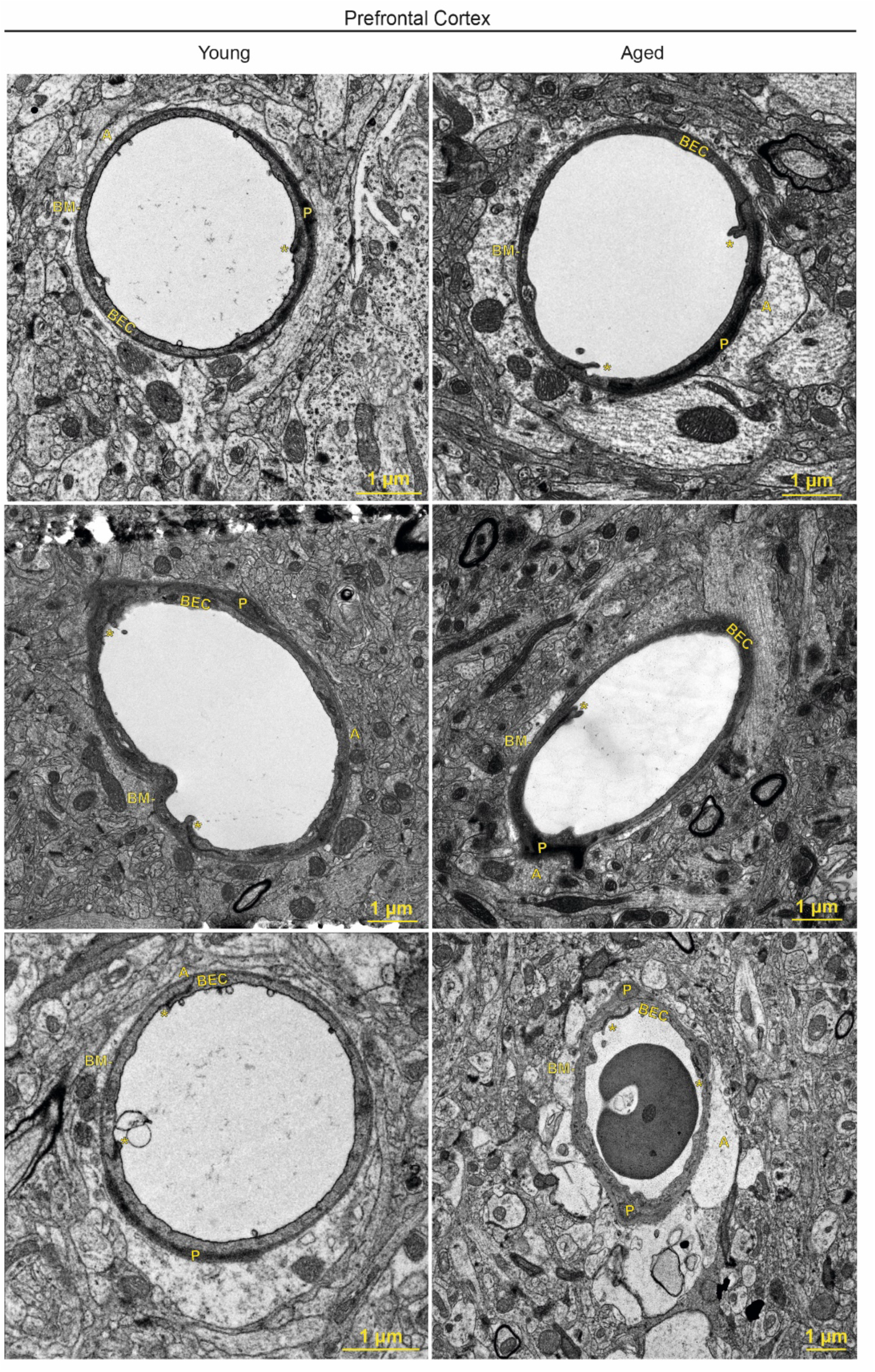
Representative images of blood vessels in the prefrontal cortex of young and aged mice. BEC: brain endothelial cell, *: tight junction, P: pericyte, BM: basement membrane, A: astrocyte.

**Supplementary Figure 5.**
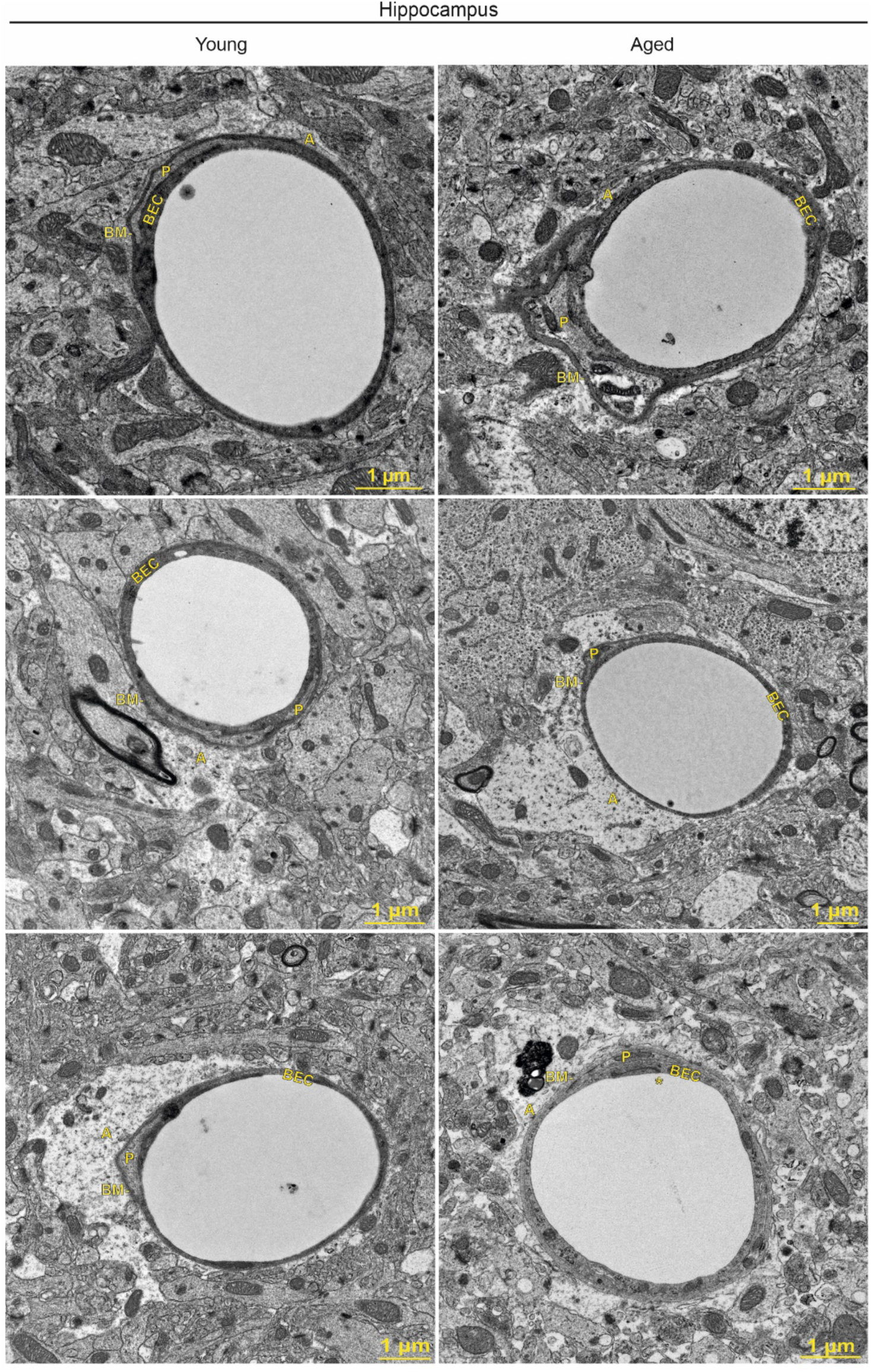
Representative images of blood vessels in the hippocampus of young and aged mice. BEC: brain endothelial cell, *: tight junction, P: pericyte, BM: basement membrane, A: astrocyte.

**Supplementary Figure 6.**
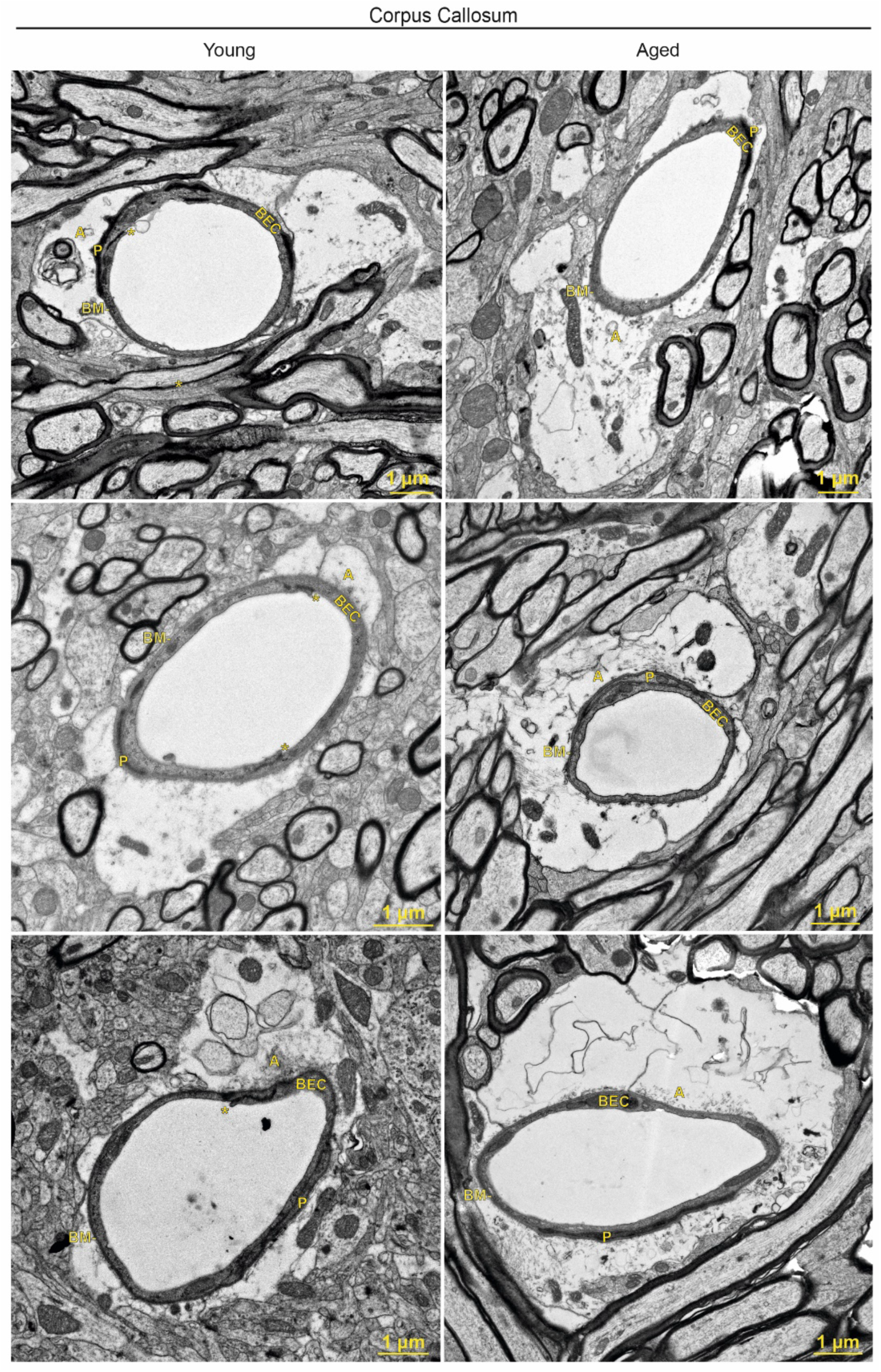
Representative images of blood vessels in the corpus callosum of young and aged mice. BEC: brain endothelial cell, *: tight junction, P: pericyte, BM: basement membrane, A: astrocyte.

